# scRepresenter: a workflow for computing, integrating and benchmarking cellular representations in single-cell transcriptomics

**DOI:** 10.64898/2026.07.15.738660

**Authors:** Guilherme Pocas, Muhammad Umar, Oliver Davis, Martin Hemberg, Andre Lamurias, András Lakatos, Muhammad Asif

## Abstract

**Motivation:** Single-cell RNA sequencing (scRNA-seq) has become an attractive tool for studying complex diseases, in which transient cell states affecting diverse cell populations characterise disease development and progression. However, due to data sparsity and disease heterogeneity analysis is often challenging. With recent advances in machine learning, two widely used approaches have emerged for learning cellular representations: large-scale foundation models and biological knowledge-guided methods. Despite their complementary strengths, there is currently no unified workflow for systematically comparing and integrating these approaches.

**Results:** Here, we present scRepresenter, an open-source workflow for computing, integrating, and validating cellular embeddings derived from foundation models and biological knowledge-guided methods in the context of complex diseases. It consists of two components: a command-line workflow that computes cellular embeddings and performs downstream analyses, and an interactive Shiny application for visualizing and comparing the computed embeddings. scRepresenter supports four categories of cellular representations: (1) expression-based, (2) knowledge-guided, (3) foundation model-derived, and (4) hybrid embeddings that combine foundation model-derived representations with knowledge-guided representations. This approach takes a cell-by-gene count matrix as input and outputs an integrated object containing the computed embeddings. Then, this object can be uploaded into our interactive Shiny application to compare different embeddings.

**Availability:** The workflow is available at https://github.com/GuilhermePocas/scRepresenter

## 1 Introduction

Single-cell RNA-sequencing (scRNA-seq) enables fine-grained profiling of cell types and cell states in development and disease(Gulati et al. 2020, Li et al. 2023, Pineda et al. 2024, Clarence et al. 2025). A typical scRNA-seq workflow involves preprocessing, low-dimensional cellular representations, and cell-type annotation, followed by downstream analyses(Wolf, Angerer, and Theis 2018, Aussel et al. 2022, Hao Y et al. 2024).

Although numerous tools exist for scRNA-seq analysis, a central challenge remains the computation of cellular representations that robustly capture the underlying biological structure from technically confounded transcriptomic data. Since cellular representations directly influence downstream analyses such as clustering, cell-type annotation, and biological interpretation, selecting an appropriate representation is a critical step in single-cell analysis. This challenge is particularly important when studying low-abundance cell populations, transcriptionally similar cell types, or disease-associated cellular states, where subtle biological signals can be easily obscured by noise and data sparsity(Dann et al. 2023).

Transformer-based foundation models have recently emerged as powerful approaches for deriving transcriptomic signatures from millions of cells(Cui et al. 2024, Hao M et al. 2024, Zeng et al. 2025). These models capture global tissue-level structure and generalise across datasets. However, their ability to encode biologically meaningful information depends on the diversity and balance of the training data. In particular, rare cell types or transient cellular states, often observed in disease-related conditions, may not have sufficient representation for training the model.

In parallel, biological knowledge-guided methods incorporate structured priors, such as protein–protein interaction (PPI) networks annotations(Ashburner et al. 2000), to improve interpretability and local biological coherence(Sheinin, Sharan, and Madi 2025). However, such methods do not explicitly model global transcriptomic hierarchy, provided by foundation models trained on millions of cells.

Despite their complementary strengths, foundation model–based and knowledge-guided approaches are typically applied independently. Combining these approaches may produce more robust cellular embeddings and as such, improve the identification of disease-relevant cell types and states. A unified workflow that enables systematic computation, comparison, and integration of these representation strategies is currently lacking.

Here, we developed scRepresenter, an open-source workflow for generating, integrating, and benchmarking multiple biologically informed cellular representations within standard single-cell analysis pipelines. scRepresenter supports expression-based embeddings, knowledge-guided embeddings, foundation model– derived embeddings, and hybrid embeddings that combine global transcriptomic context with structured biological priors. The package includes a command-line workflow for embedding computation and downstream evaluation, together with an interactive Shiny application for comparative visualisation and exploration of the resulting representations.

To demonstrate the utility of the embeddings computed by scRepresenter, we applied it to a complex disease dataset. Downstream analysis showed that combining foundation model embeddings with knowledge-guided embeddings better recapitulated biology, preserved reference cell-type structure and lineage trajectories in a brain organoid case study.

## 2. Workflow implementation and architecture

scRepresenter consists of two main components: a command-line workflow for computing cellular embeddings and an interactive Shiny application for comparing, exploring, and visualizing the resulting representations. Figure 1A shows the architecture of the command-line workflow, which is implemented in Python and can be installed using Conda or run through a Docker image. It takes a processed scRNA-seq AnnData object as input, along with parameters for model fine tuning. Then it outputs individual embeddings for each representation strategy (detailed instructions for installation, preprocessing, input and output formats are provided in the Supplementary Material).

**Figure 1.**
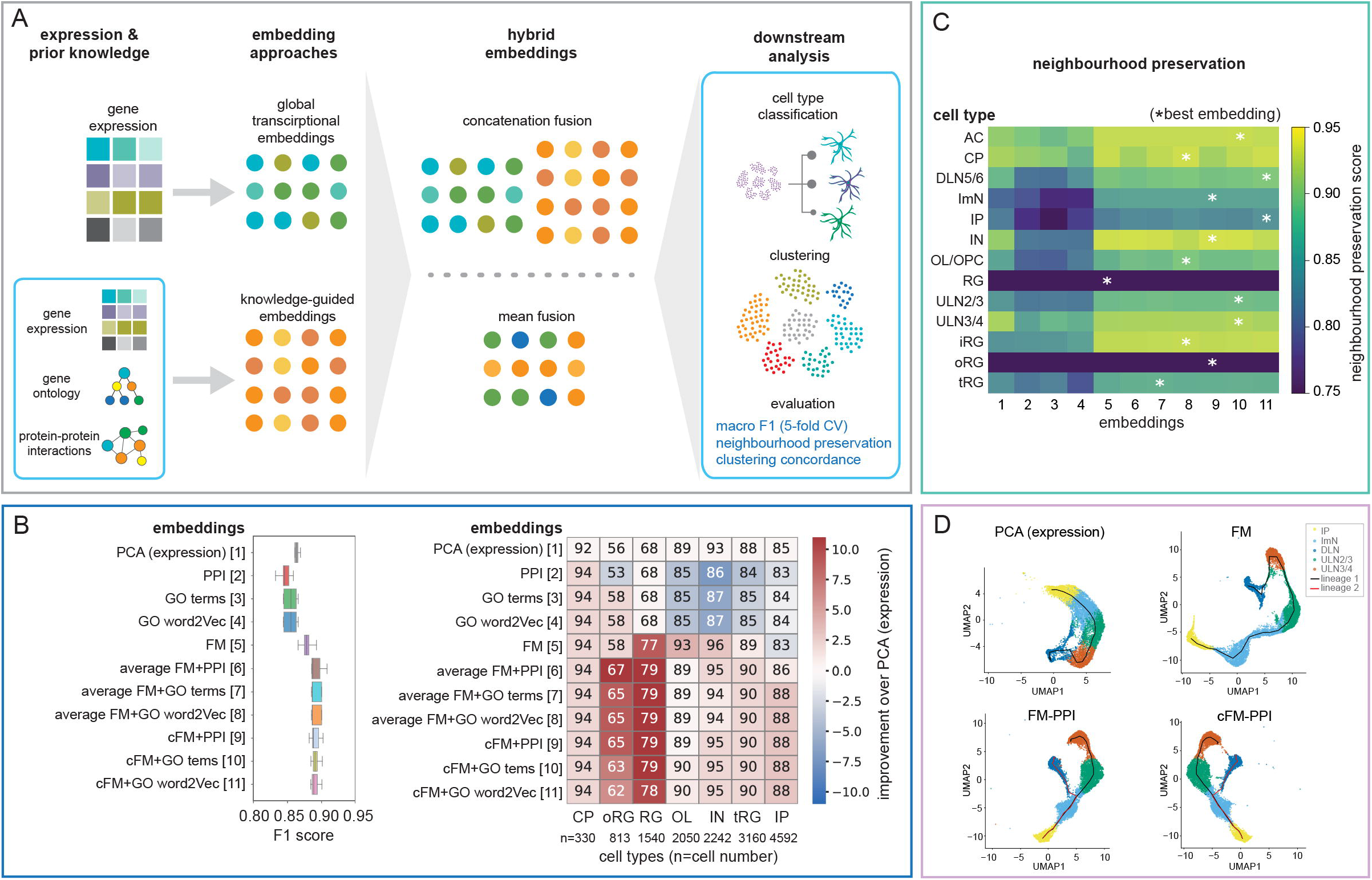
scRepresenter architecture and benchmarking using human neurodegenerative disease datasets. **(A)** Overview of the scRepresenter modular architecture of the workflow (from left to right), illustrating data pre-processing, generation of foundation model embeddings, integration of structured biological knowledge, construction of hybrid embeddings, and downstream supervised and unsupervised analyses. **(B)** Supervised evaluations strategies used by scRepresenter. Boxplots (left) summarising cell-type classification performance across stratified five-fold cross-validation for each embedding strategy, evaluated using the macro F1-score. Heatmap (right) showing the cell type–specific classification accuracy for each embedding strategy. Values represent the proportion of correctly classified cells per cell type; n=number of cells. The colour gradient represents the percentage values of improvement relative to the baseline PCA (expression) performance. **(C)** Unsupervised evaluations strategies employed by scRepresenter. Heatmap of cell type–specific neighbourhood preservation scores for each embedding. Neighbourhood preservation was defined as the fraction of k-nearest neighbours sharing the same reference cell-type label. Asterisks indicate the highest score per cell type. **(D)** Pseudotime trajectory analysis. UMAPs showing cellular lineages captured by selected embeddings.

The command-line workflow supports expression- and foundation model-derived, knowledge-guided, and hybrid cellular representations. For foundation model-derived embeddings, scRepresenter incorporates scGPT, a transformer-based model trained on more than 33 million single-cell transcriptomes(Cui et al. 2024). We selected the scGPT pretrained checkpoint for brain cells. However, scRepresenter is open source and modular, allowing users to modify the framework and incorporate other foundation models as needed. For knowledge-guided embeddings, it includes an optimised implementation of scNET, which integrates gene expression data with biological graphs derived from PPI networks(Sheinin, Sharan and Madi 2025). scRepresenter currently supports PPI networks and added two additional gene ontology (GO) derived gene similarity networks as prior biological knowledge, which were missing in scNET.

Both scGPT and scNET output a vector of 512 dimensions. Hybrid embeddings are generated by combining scGPT and scNET representations using either element-wise averaging or feature concatenation. Embeddings are normalised before averaging them. For concatenation, standardized embedding vectors are joined and optionally dimensionally reduced using PCA before downstream analysis (details on model fine-tuning, biological network preparation, embedding generation and integration are provided in the Supplementary Material).

For downstream evaluation, scRepresenter interfaces with Scanpy for unsupervised analyses and implements supervised cell-type classification using a neural network classifier with stratified five-fold cross-validation. Supervised evaluation reports macro F1-score and cell-type-specific prediction accuracy, enabling users to assess how well each representation supports cell-type classification. Unsupervised evaluation compares the preservation of reference cell-type structure using neighbourhood preservation and pseudotime trajectory analysis to understand cellular differentiation trajectory for each embedding. Together, these outputs allow users to identify which representation strategy best preserves biologically meaningful structure in a given dataset.

The combined output object generated by the command-line workflow can be loaded directly into the R Shiny application. The application provides an interactive graphical interface for comparing embeddings, including low-dimensional projections, cell-type-specific performance heatmaps, prediction summaries, misclassification patterns, and Sankey plots showing label correspondence across embedding strategies. All plots can be exported for downstream reporting and publication (the Shiny application is available from the scRepresenter GitHub repository, with a tutorial).

## 3. Application to ground truth and complex scRNA-seq data

First, scRepresenter was evaluated on peripheral blood mononuclear cell (PBMC), glioblastoma mouse (GBM), and pancreas baron datasets. These datasets have been widely used as ground-truth reference to benchmark the performance of single cell analysis methods. Embeddings were computed separately for each dataset using the command-line module. For these ground-truth reference datasets, embedding performance was assessed by cell-type classification using a neural network classifier, with the F1 score used as the main evaluation metric. The results showed that hybrid embeddings more accurately predicted cell-type labels across benchmark datasets (details on the datasets and classification performance are provided in the GitHub repository and Supplementary Material). To demonstrate the utility of scRepresenter in a challenging scenario, it was run on a complex scRNA-seq dataset from human brain organoid neurodegeneration model, 3D tissues generated from patient-derived stem cells harboring the amyotrophic lateral sclerosis (ALS)-causing C9ORF72 mutation(Szebényi et al. 2021). This disease-relevant system provides a challenging setting for representation benchmarking because cell types vary greatly both in abundance and transcriptional signatures in vitro (the Supplementary Material provides details of the dataset, preprocessing, model training, and evaluation strategy).

For each representation generated by scRepresenter, supervised and unsupervised evaluations were performed. For supervised evaluations, embeddings were projected to a lower-dimensional space using PCA and a neural network with stratified five-fold cross-validation was trained using the cell labels from Szebényi *et al*. (Szebényi *et al*. 2021 as reference. Macro F1-score was selected as the primary metric because it summarizes classification performance across cell types and is less dominated by abundant populations. In this case study, hybrid representations achieved the highest macro F1-scores (Figure 1B). Improvements were most pronounced for low-abundance and transcriptionally similar populations. For example, correct identification of outer radial glial cells increased to 67% with hybrid embeddings, compared with 58% using either foundation model-derived or knowledge-guided embeddings alone. Similar improvements were observed for radial glial cells, indicating that hybrid embeddings could have improved resolution of closely related progenitor populations.

We next evaluated whether the embeddings preserved intrinsic cell-type structure. Local neighbourhood preservation was assessed using k-nearest-neighbour graphs constructed with cosine similarity. For each cell, the neighbourhood preservation score was defined as the proportion of nearest neighbors sharing the same reference cell-type label. Hybrid embeddings showed higher neighborhood preservation across multiple cell types (Figure 1C).

Having identified technical improvements using combined embedding approach, we next asked whether the new embeddings better captured the underlying biology. To do this, we used Slingshot to infer the neurodevelopmental trajectories of intermediate progenitor cells, which normally differentiate into either upper or deep layer neuronal lineages. This was effectively captured in hybrid embeddings, but not in the PCA-based and foundation-model derived embeddings (Figure 1D). Details on the pseudotime trajectory analysis can be found in the Supplementary Material.

Together, these results demonstrate that hybrid embeddings provided the strongest preservation of reference labels, local neighbourhood structure, and inferred differentiation trajectories. This case study also illustrates how scRepresenter can be used to compare embedding strategies in complex scRNA-seq datasets.

## 4. Conclusions

scRepresenter provides an open-source workflow for computing, integrating, visualising, and benchmarking cellular representations in scRNA-seq data. By combining command-line embedding generation with an interactive Shiny interface, it enables users to compare expression-based, knowledge-guided, foundation-model-derived, and hybrid representations within a unified framework. In the ALS brain organoid case study, hybrid embeddings improved the preservation of reference cell-type structure, local neighbourhood organization, and inferred differentiation trajectories. However, our data also reflects that representation performance is dataset-dependent, and the optimal strategy may vary across biological systems, tissue types, and disease contexts. scRepresenter is therefore designed to support systematic evaluation and selection of appropriate cellular embeddings for individual datasets. Our workflow provides a grounding for further improvements, including the employment of additional representation methods, broader benchmark datasets, and upgraded support for computationally efficient foundation-model workflows.

## Abbreviations

AC: astrocyte/astroglia
CP: choroid plexus
DLN: deep-layer 5/6 excitatory neurons
IE: immature excitatory neurons
IP: intermediate progenitors
IN: interneurons
OLG/OPC: oligodendrocytes and oligodendrocyte precursor cells
RG: radial glia
ULN2/3: upper-layer 2/3 excitatory neurons
ULN3/4: upper-layer 3/4 excitatory neurons
iRG: intermediate radial glia
oRG: outer radial glia
tRG: truncated radial glia
PPI: protein-protein interaction network
FM: foundation model
cFM: concatenated foundation model
GO: gene ontology

## Author contributions

Concept and project leads: M.A., A. Lakatos, computational design: M.A., G.P., M.U., A.L., data analysis: G.P., M.A., data interpretation: G.P., M.U., M.A., M.H., A.L., A. Lakatos, writing – first draft: M.A. with contribution from all authors., manuscript editing: A. Lakatos, M.H., A.L.

