## Supplementary Material for "scRepresenter: a workflow for computing, integrating and benchmarking cellular representations in single-cell transcriptomics"

**scRepresenter -** Supplementary Material

scRepresenter is an open-source workflow for computing biologically informed cellular representations from single-cell RNA sequencing (scRNA-seq) data. The workflow computes expression-derived, foundation model-derived, biological knowledge-guided, and hybrid embeddings (combining the foundation model-derived and biological knowledge-guided embeddings) and outputs a common object containing all embeddings. This enables users to compare how different representation strategies affect cell-type classification, neighbourhood structure, clustering, and biological interpretability. By combining large-scale foundation models with biological priors from other relevant sources, it is possible to gather insights from millions of cell expression profiles while also capturing more local, biologically relevant associations that might be hidden in the scRNA-seq data from the current experiment. This complementary approach was the basis for developing the scRepresenter pipeline.

### 1. scRepresenter architecture

scRepresenter consists of a command line module and a shiny R interactive app which can be installed and run easily.

#### 1.1. Command line module

This module is comprised of two main components, one focused on a data-driven approach, that takes advantage of established foundation model architectures, and another responsible for the integration of relevant biological knowledge.

##### 1.1.1. Data-driven foundation model-based approach

Complex foundation model architectures like GeneFormer (Theodoris et al. 2023), scFoundation (Hao et al. 2024) and scGPT (Cui et al. 2024) are able to find biological insights from millions of cells. Selecting the foundation model is challenging because available models differ in the datasets, biological contexts, and training strategies used during pretraining. Since one of the core objectives of scRepresenter was to evaluate cellular representations in complex diseases, including neurodegenerative diseases, we selected the scGPT pretrained checkpoint for brain cells. However, scRepresenter is open source and modular, allowing users to modify the framework and incorporate other foundation models as needed.

The scGPT foundation model itself is a large transformer architecture that uses a masked language modelling (MLM) learning strategy to achieve better results in a variety of tasks. It also offers a range of pretrained checkpoints, that can be seen in Table 1.

Table 1. Pretrained checkpoints for the scGPT foundation model.

| **Checkpoint same** | **Pretraining**  **dataset size** | **Description** |
| --- | --- | --- |
| Whole-human | 33,000,000 | Largest, general-purpose model |
| Continual pretrained | - | Adjusts the Whole-human model for zero-shot tasks (no fine-tuning) |
| Brain | 13,200,000 | Pretrained on brain cells |
| Blood | 10,300,000 | Pretrained on blood and bone marrow cells |
| Heart | 1,800,000 | Pretrained on heart cells |
| Lung | 2,100,000 | Pretrained on lung cells |
| Kidney | 814,000 | Pretrained on kidney cells |
| Pan-cancer | 5,700,000 | Pretrained on several types of cancer cells |

##### 1.1.2. Biologically enhanced learning

##### The second component of scRepresenter’s command-line module is responsible for incorporating biological priors into the exclusively expression-based embeddings. The motivation for this decision is that integrating additional knowledge of biological relevance, such as gene programs or pathways, has been shown to improve the robustness of downstream analysis. This becomes even more relevant in the context of scRNA-seq datasets, which are inherently noisy and sparse and may benefit greatly from more structured data sources, such as hierarchical databases or ontologies.

For this purpose, we chose scNET, a network-based hybrid architecture that offers several advantages over other techniques for incorporating biological knowledge. First off, since scNET is an unsupervised method, there is no concern for data leakage when combining its output with other models, which is especially important for our purposes. Additionally, this model employs a gene similarity network from which it collects biologically relevant associations, meaning it allows for a lot of flexibility in how this network is constructed and on which sources are used to do so. Table 2 lists the biological networks currently supported by scRepresenter.

Table 2. scRepresenter’s biological networks.

| **Biological context** | **Source** | **Genes N.** | **Associations N.** |
| --- | --- | --- | --- |
| Protein-protein interactions | ANAT 3.0 | 19,043 | 544,455 |
| Gene  function | Gene  Ontology | 44,598 | 784,711 |

###### 1.1.2.1. Gene Similarity Networks

###### The similarity network is essentially an undirected gene graph, where each node represents a single gene, and each edge represents an association between genes, with a similarity weight calculated with respect to a specific biological context. Both the choice of knowledge source from which we extract the biological similarity between genes and the way the network is constructed are crucial to ensuring the model functions properly. Originally, scNET employed a network based on Protein-Protein Interactions (PPIs); however, we also considered associations from the Gene Ontology (GO), with Table 2 showing how the two sources compare.

Having established sources from where we can collect relevant biological information is important; however, how this data is organised into a machine-learning-readable network is also crucial for the model to achieve a good performance. With this goal, we devised two different methods to construct a weighted gene graph from our knowledge sources, for a given scRNA-seq dataset.

###### Common terms method

The first method focuses on the shared entries among genes to determine how closely related they are, meaning it focuses only on GO associations and does not directly consider the Ontology’s structure. This technique is illustrated in Figure 1. First, the top 3000 most variable genes in a given scRNA-seq dataset are selected, and each gene is mapped to its corresponding list of GO terms in the ontology; genes with no associated terms are excluded. From the remaining genes, we compute all possible gene pairs and count the number of terms they have in common across their corresponding term lists, creating a gene similarity matrix from these counts. Finally, for each gene, we select only the top 100 pairs with the most terms in common to ensure the network remains a reasonable size. After arriving at the final values, the logarithm transformation and min-max normalisation are applied to prevent outliers from skewing the network's balance.

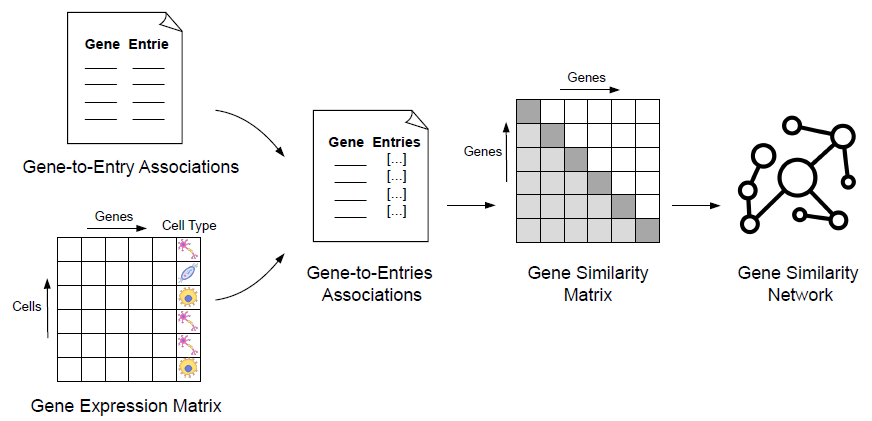

**Figure 1.** The common terms method, where the similarity weight between genes is based on how many terms they have in common in a given biological knowledge source.

With 𝑇(𝑔) representing the set of terms associated with gene 𝑔, from one of the knowledge sources, the similarity between two genes 𝑔𝑖 and 𝑔𝑗 can be defined as:

$$S\left( g_{i},g_{j} \right)=\frac{log\left( 1+\left| T\left( g_{i} \right)\cap T\left( g_{j} \right) \right| \right)-s_{min}}{s_{max}-s_{min}}$$

Where:

- $\left| T\left( g_{i} \right)\cap T\left( g_{j} \right) \right|$ is the number of shared terms between the two genes.
- log(1 + 𝑥) is applied to reduce the effect of large overlaps.
- $s_{min}$ and $s_{max}$ are the minimum and maximum used for min-max normalization.

This results in a similarity score 𝑆(𝑔𝑖 , 𝑔𝑗) ∈ [0, 1], that will be used to weight the edges of the resulting gene similarity graph.

###### GO Embeddings Method

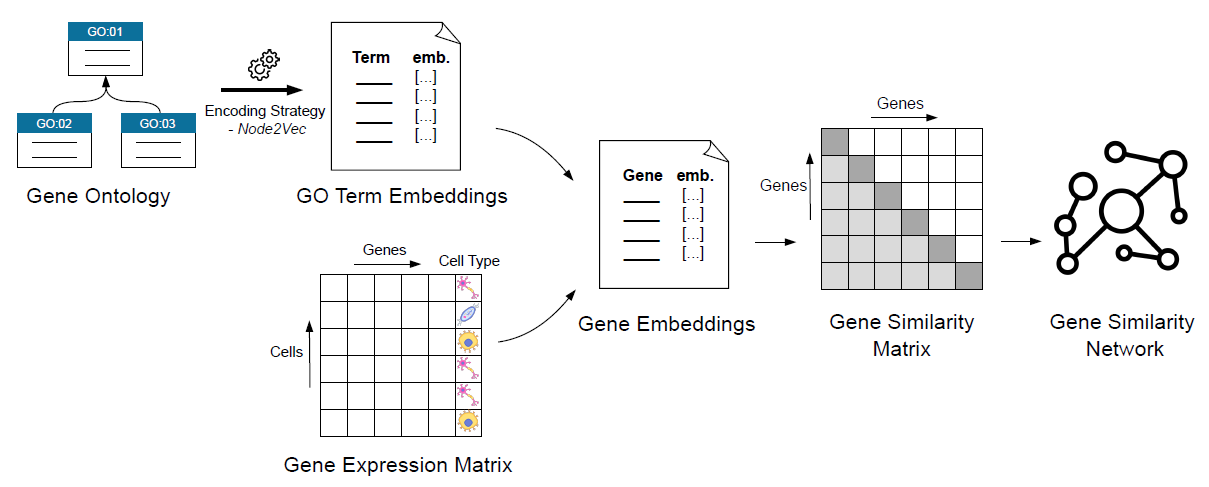
This method aims to not only capture the relationship between genes based on their associated terms, but also on their hierarchical position in the Gene Ontology. The pipeline for this method is illustrated in Figure 2.

**Figure 2.** The GO embeddings methos, where the Gene Ontology is encoded into a set of embeddings, that are used to calculate the similarity weight between genes.

In order to capture the structure of the ontology, this method applies an additional embedding algorithm, Node2Vec (Grover and Leskovec 2016) (based on the NLP method Word2Vec (Mikolov et al. 2013)), to the GO so that each GO term will have a corresponding numerical representation in the same shared latent space, capturing its position in the ontology’s hierarchy. With these embeddings, it’s possible to then calculate a numerical representation for each gene, by calculating the element-wise average of the embeddings of every associated GO term. So, for a gene 𝑔 and its term set 𝑇(𝑔), where each term 𝑡 has an assigned term embedding $e_{t}$ , the corresponding gene embedding $e_{g}$ is defined as:

$$e_{g}=\frac{1}{T\left( g \right)\vee\sum_{t\in T\left( g \right)} e_{t}}$$

After having calculated a set of gene embeddings, a similarity metric for each gene pair can then be calculated. So, for the top 3000 highest variable genes, and for all possible gene pairs, the cosine similarity between both gene embeddings is calculated and used to indicate the degree to which they are related. So, for a given pair of genes $g_{i}$ and $g_{j}$, and their corresponding embeddings $e_{g_{i}}$ and $e_{g_{j}}$, the similarity weight will be:

$$S\left( e_{g_{i}},e_{g_{j}} \right)=\frac{e_{g_{i}}.e_{g_{j}}}{\vee e_{g_{i}}\vee\vee e_{g_{j}}\vee}$$

The resulting score $S\left( e_{g_{i}},e_{g_{j}} \right)\in\left[ 0,1 \right]$, will again be used to weight the edges of the resulting similarity graph.

#### 1.1.3. Hybrid Embeddings

With the output of both models each cell of a given scRNA-seq dataset will have two vector representations, one focused on the biological priors present in the gene network, and another that more closely represents the expression profile of that cell. We opted for two simple hybridization methods to prove the combination of both models resulted in simple and measurable improvements.

The first technique performs the element-wise average of each cell’s embeddings and calculates the resulting mean hybrid representation. Formally, for a given cell’s embeddings $e^{scNET}$ and $e^{scGPT}$ with a dimension size of $d$, the hybrid embedding $e^{avg}$ will also have $d$ dimensions that can be defined by:

$$e_{i}^{avg}=\frac{e_{i}^{scNET}+e_{i}^{scGPT}}{2},i=1,\ldots,d$$

The second technique is similar, however both embeddings are now concatenated, leading to a single numerical vector with twice the number of elements. Formally, for the same embeddings as before, $e^{scNET}$ and $e^{scGPT}$, with a dimension size of $d$, the concatenate embedding $e^{con}$ can be defined by:

$$e^{con}=\left( e_{1}^{scNET},\ldots,e_{d}^{scNET},e_{1}^{scGPT},\ldots,e_{d}^{scGPT} \right)$$

In order to avoid excessive dimensions, the PCA dimensionality reduction technique is then applied to reduce the dimensions back to their original size.

#### 2. Case studies datasets and analysis

To demonstrate the applicability of scRepresenter, we applied it to multiple ground-truth datasets. The computed embeddings were evaluated for cell-type prediction using the reference labels provided with each dataset. For this evaluation, we trained a neural network classifier.

Overall, we considered the following four setups:

- **Expression-based classification:** Thi**s** setup represents the baseline for the standard classification of scRNA-seq data. It uses embeddings calculated from the PCA-reduced expression profiles of each cell, allowing for comparisons to be made against more complex methods.
- **Knowledge-based classification**: This setup utilizes primarily the scNET model to produce a set of cell representations, in order to test how biological priors can impact the classification of cell type. It also allows for the analysis of different biological contexts and their effect on classification performance, by using the three differently build networks explained previously.
- **Foundation model-based classification:** This method focuses on the scGPT foundation model, that computes cell representations from single cell data. It has complex architecture and has been pretrained on millions of cells.
- **Hybrid classification**: Finally, this is the scRepresenter workflow in its entirety, combining the previous two methods into a single set of cell-level embeddings. By comparing its performance with each of the previous methods, we can understand the benefits and potential limitations of our complementary approach.

The classification process consists of a fivefold cross-validation loop, in which a neural network-based classifier architecture is fine-tuned on 4 folds of a scRNA-seq dataset before predicting the labels of the remaining test fold. The architecture consists of a standard three-layer supervised architecture, with the first layer projecting the initial 512 embedding dimensions into 128, followed by another hidden layer projecting them into 64, and the final layer mapping them to the number of cell types present in that dataset. At each step, the GELU activation function is also applied.

Additionally, since some of the representation learning methods used for embedding creation are supervised, mainly scGPT, and by consequence scRepresenter, there was a risk of data leakage when producing embeddings for the entire dataset. As such, in the embedding process, there is a train/test split, with 60% of the smaller datasets (<10,000 cells) or 80% of the larger datasets (>10,000 cells) used exclusively to train the model. The remaining cells have no impact on model training and are reserved for cross-validation classification to avoid potential performance inflation.

The above mentioned experimental setups were applied to multiple datasets (Table 3)

Table 3. Datasets used for the case studies.

| **Dataset** | **Cell N.** | **Gene N.** | **Cell types N.** |
| --- | --- | --- | --- |
| PBMC | 2,638 | 32,738 | 8 |
| GBM | 7,273 | 18,531 | 8 |
| Baron | 8,569 | 17,499 | 14 |
| ALS brain organoids | 75,497 | 3,000 | 13 |

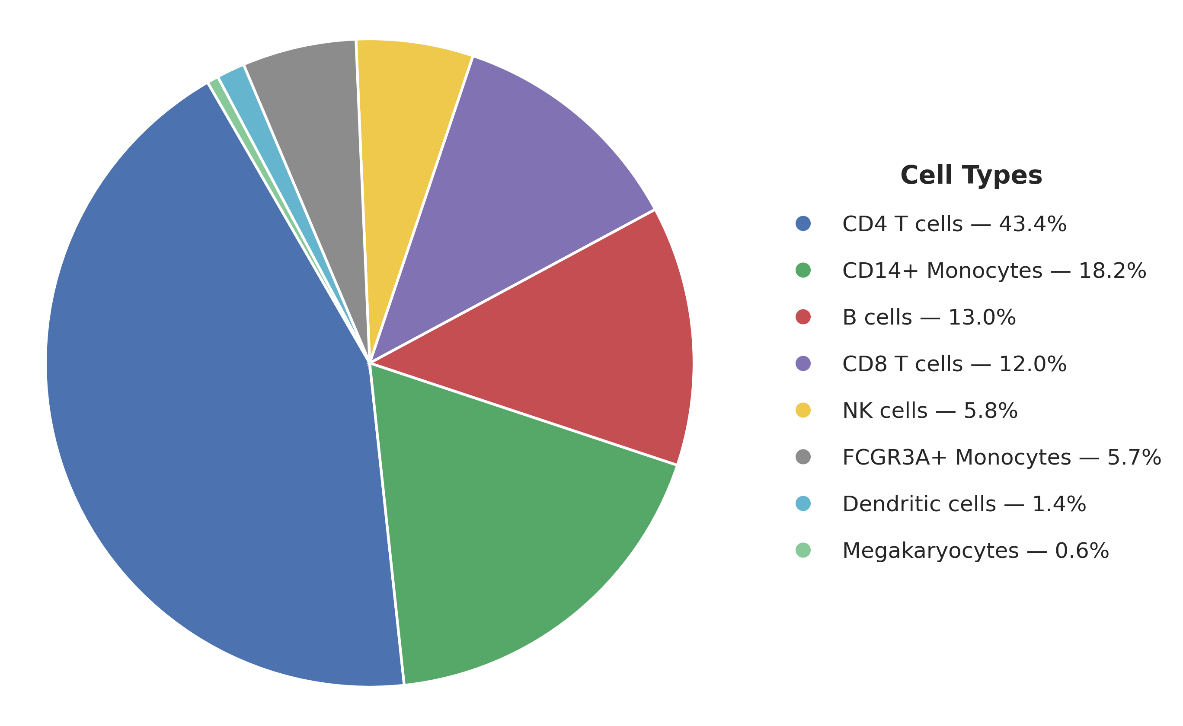

#### 2.1 PBMC Dataset

**Figure 3.** Cell type distribution for the PBMC dataset.

This peripheral blood mononuclear cell (PBMC) dataset^[1]^ offers a relatively small cell count, which makes it easier to over or undertrain models when applying machine learning architectures. Another aspect to consider is that almost half of the dataset is composed of CD4 T cells, with the remaining cell types making up the other half. Such a large disparity also means that models may overly focus on CD4 T cells at the cost of performance when predicting less common cell types.

##### 2.1.1 PBMC classification performance

The average classification performance on the PBMC dataset across all setups is shown in Table 4. This includes the four setups mentioned previously, which, when considering the three available gene networks and the two possible hybridisation methods, come down to a total of eleven possible testing setups.

Table 4. Classification performance for the PBMC dataset across all the considered setups. The values shown for each of the four metrics of accuracy, precision, recall and F1-score is a mean of the five corresponding values obtained for each fold of the fivefold cross validation classification setup.

| **Method** |  | **Accuracy** | **Precision** | **Recall** | **F1-score** |
| --- | --- | --- | --- | --- | --- |
| PCA for expression |  | 0.9086 | 0.8966 | 0.7730 | 0.7917 |
| PPI |  | 0.9356 | 0.9385 | 0.9040 | 0.9068 |
| GO common-terms |  | 0.9366 | 0.9446 | 0.8902 | 0.9008 |
| GO word2vec |  | 0.9375 | 0.9459 | 0.9018 | 0.9091 |
| FM |  | 0.9403 | 0.9335 | 0.9147 | 0.9079 |
| Average FM and PPI |  | 0.9460 | 0.9506 | 0.9241 | 0.9238 |
| Average FM and GO common-terms |  | 0.9451 | 0.9471 | 0.9185 | 0.9190 |
| Average FM and GO word2vec |  | 0.9423 | 0.9480 | 0.9200 | 0.9201 |
| cFM and PPI |  | 0.9508 | 0.9559 | 0.9259 | 0.9268 |
| cFM and GO common-terms |  | 0.9489 | 0.9496 | 0.9267 | 0.9245 |
| cFM and GO word2vec |  | 0.9508 | 0.9542 | 0.9247 | 0.9253 |

Recall and F1-score for cell type classification were lower than the accuracy and precision, probably due to the presence of rare cell types in the dataset. The baseline PCA setup especially struggles with this, achieving by far the lowest recall and F1-score among all tested conditions. Both the foundation model and the biologically enhanced setups significantly improve on the baseline, reaching around a 90% F1-score, indicating that either approach is better than the simple PCA of the expression values, as expected. Still, all hybrid conditions show a much greater improvement, with the concatenate aggregation method having the highest values overall.

2.2 GBM Dataset

The Glioblastoma Mouse (GBM) dataset (Yeini et al. 2021) has a relatively balanced distribution of cells types. This property makes it well suited for classification, as even simpler models have plenty of training examples of the least common cells, leading to more accurate predictions.

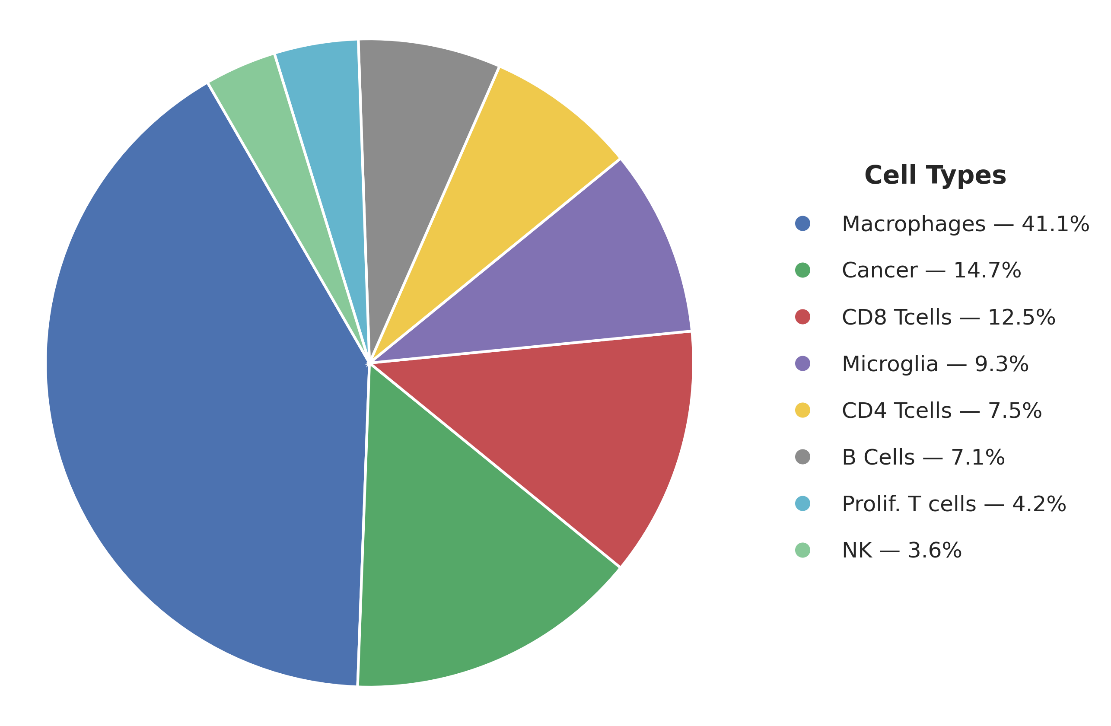

**Figure 4.** Cell type distribution for the GBM dataset.

##### 2.2.1 GBM classification performance

As expected, the performance metrics are noticeably higher, even including the baseline PCA-only test setup. Additionally, the recall and the F1-score are also closer to the accuracy and precision values, indicating that the cell type predictions are much more balanced, consequence of the balanced cell type ratios of this dataset (Table 5).

In this case, the introduction of biological priors shows no major improvement from the baseline setup, with both showing a comparable performance. The foundation model-based setup, however, shows a significant increase across all metrics, showing that the complex architecture of this method might be especially relevant in this dataset. Finally, while not as pronounced as in other datasets, there is a similar increase in performance across most hybrid embeddings, suggesting there is still a benefit to using the hybrid approach in well-balanced datasets.

Table 5. Classification performance for the GBM dataset across all the considered setups. The values shown for each of the four metrics of accuracy, precision, recall and F1-score is a mean of the five corresponding values obtained for each fold of the fivefold cross validation classification setup.

| **Method** |  | **Accuracy** | **Precision** | **Recall** | **F1-score** |
| --- | --- | --- | --- | --- | --- |
| PCA for expression |  | 0.9527 | 0.9464 | 0.9300 | 0.9375 |
| PPI |  | 0.9601 | 0.9447 | 0.9374 | 0.9399 |
| GO common-terms |  | 0.9581 | 0.9481 | 0.9340 | 0.9393 |
| GO word2vec |  | 0.9563 | 0.9364 | 0.9287 | 0.9313 |
| FM |  | 0.9618 | 0.9471 | 0.9423 | 0.9430 |
| Average FM and PPI |  | 0.9691 | 0.9583 | 0.9550 | 0.9558 |
| Average FM and GO common-terms |  | 0.9663 | 0.9555 | 0.9475 | 0.9496 |
| Average FM and GO word2vec |  | 0.9649 | 0.9523 | 0.9448 | 0.9468 |
| cFM and PPI |  | 0.9684 | 0.9581 | 0.9511 | 0.9538 |
| cFM and GO common-terms |  | 0.9708 | 0.9612 | 0.9519 | 0.9548 |
| cFM and GO word2vec |  | 0.9691 | 0.9561 | 0.9541 | 0.9536 |

#### 2.3 Baron Dataset

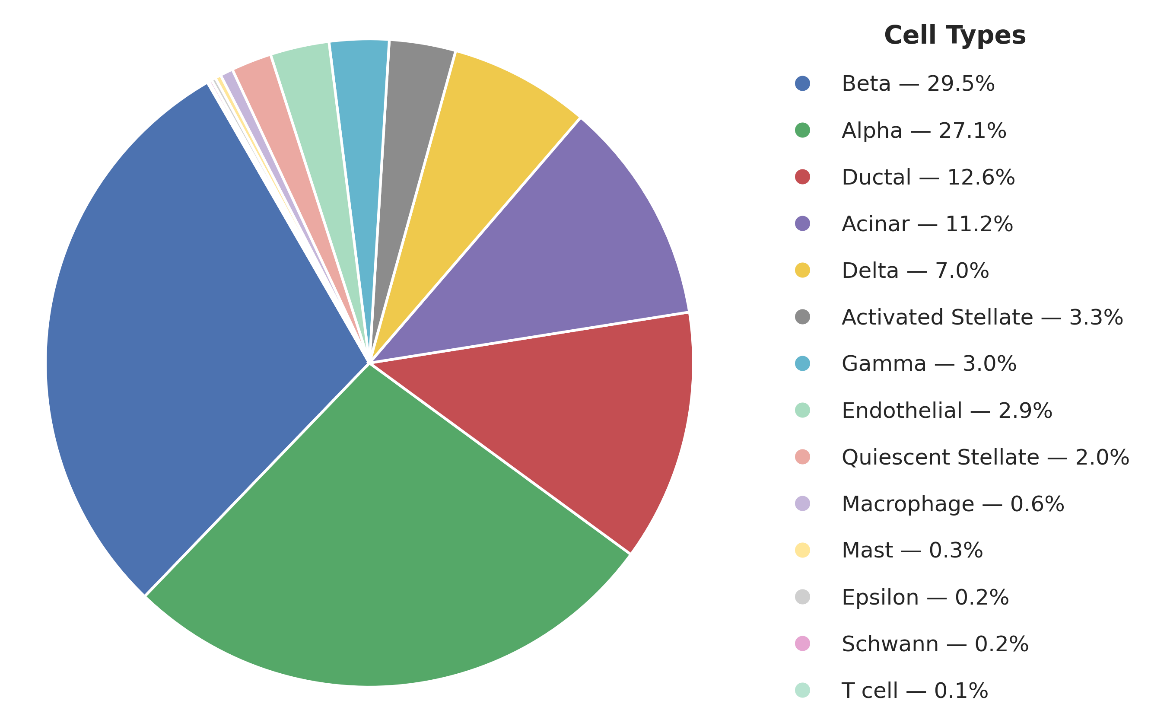
The Baron dataset (Abdelaal et al. 2019) offers unbalanced testing conditions, with the five least common cell types making up only around 1% of the total number of cells. Because of this, it is expected that most setups will struggle with balanced set of predictions.

**Figure 5.** Cell type distribution for the Baron dataset.

##### 2.3.1 Baron classification performance

The performance metrics reflect the dataset’s properties: most setups achieve very high accuracies due to many correct predictions on simpler, common cells, but a comparatively low recall and F1-score. This is most noticeable in the baseline and biologically enhanced testing setups, with the introduction of biological priors also reducing classification precision.

Table 6. Classification performance for the Baron dataset across all the considered setups. The values shown for each of the four metrics of accuracy, precision, recall and F1-score is a mean of the five corresponding values obtained for each fold of the fivefold cross validation classification setup.

| **Method** |  | **Accuracy** | **Precision** | **Recall** | **F1-score** |
| --- | --- | --- | --- | --- | --- |
| PCA for expression |  | 0.9775 | 0.9752 | 0.7847 | 0.7941 |
| PPI |  | 0.9746 | 0.9376 | 0.8027 | 0.7999 |
| GO common-terms |  | 0.9737 | 0.9285 | 0.7892 | 0.7816 |
| GO word2vec |  | 0.9755 | 0.9506 | 0.7941 | 0.7874 |
| FM |  | 0.9828 | 0.9837 | 0.9483 | 0.9500 |
| Average FM and PPI |  | 0.9869 | 0.9868 | 0.9668 | 0.9681 |
| Average FM and GO common-terms |  | 0.9851 | 0.9865 | 0.9643 | 0.9667 |
| Average FM and GO word2vec |  | 0.9877 | 0.9889 | 0.9676 | 0.9697 |
| cFM and PPI |  | 0.9863 | 0.9868 | 0.9498 | 0.9525 |
| cFM and GO common-terms |  | 0.9854 | 0.9875 | 0.9659 | 0.9681 |
| cFM and GO word2vec |  | 0.9863 | 0.9878 | 0.9655 | 0.9680 |

The foundation model-based setup is again where the highest performance increase happens, with comparable values for accuracy and precision, but a much higher recall and F1-score than the previous setups. Hybrid embeddings showed higher recall and F1-score even more than the foundation model by itself. This is evidence that the addition of biological priors is enabling the data-driven approach to better discern between some key rare cells, resulting in more balanced predictions.

#### 2.4 Amyotrophic lateral sclerosis brain organoid dataset

The amyotrophic lateral sclerosis (ALS) dataset (Szebényi et al. 2021) represent lab-grown brain organoids harbouring the C9ORF72 ALS-causing mutation or its isogenic gene corrected variant. It introduces several new challenges for the classification goal, making it an ideal case-study. First, it has a much higher cell count than previous datasets, with many different cell types and population sizes. Additionally, due to the nature of brain organoid-based data, there are many differentiating cells that may display expression signatures of more than one cell type, making these cells more ambiguous and harder to classify.

**2.4.1 ALS Brain classification performance**

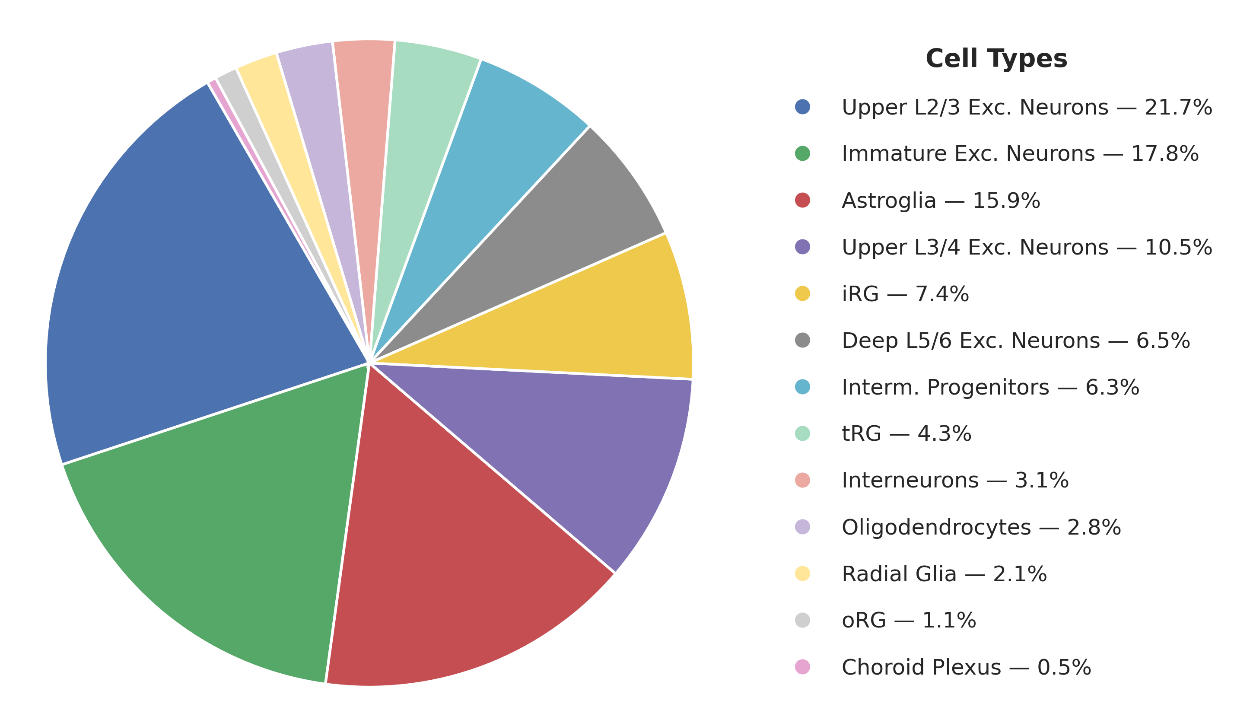

**Figure 6.** Cell type distribution for the ALS brain organoids dataset.

Compared with ground-truth datasets, the classification performance on this dataset is low across all metrics. This suggests that, unlike in previous cases where low recall and F1-score were consequences of rare, complex cell types, even accuracy and precision are lower, pointing to the complexity of the cells analysed.

Table 7. Classification performance for the ALS Brain dataset across all the considered setups. The values shown for each of the four metrics of accuracy, precision, recall and F1-score is a mean of the five corresponding values obtained for each fold of the fivefold cross validation classification setup.

| **Method** |  | **Accuracy** | **Precision** | **Recall** | **F1-score** |
| --- | --- | --- | --- | --- | --- |
| PCA for expression |  | 0.8942 | 0.8692 | 0.8604 | 0.8639 |
| PPI |  | 0.8797 | 0.8548 | 0.8407 | 0.8465 |
| GO common-terms |  | 0.8848 | 0.8634 | 0.8487 | 0.8544 |
| GO word2vec |  | 0.8848 | 0.8634 | 0.8487 | 0.8544 |
| FM |  | 0.9022 | 0.8846 | 0.8767 | 0.8784 |
| Average FM and PPI |  | 0.9152 | 0.8957 | 0.8916 | 0.8927 |
| Average FM and GO common-terms |  | 0.9156 | 0.8946 | 0.8903 | 0.8915 |
| Average FM and GO word2vec |  | 0.9156 | 0.8946 | 0.8903 | 0.8915 |
| cFM and PPI |  | 0.9157 | 0.8928 | 0.8907 | 0.8909 |
| cFM and GO common-terms |  | 0.9155 | 0.8953 | 0.8891 | 0.8914 |
| cFM and GO word2vec |  | 0.9154 | 0.8944 | 0.8885 | 0.8906 |

Consistent with the ground truth datasets, hybrid embeddings showed improved F1-score for cell type classification.

**2.5 Pseudotime and trajectory analysis of scRepresenter embeddings to explore the cellular differentiation**

Pseudotime trajectory analysis has been extensively used in single-cell transcriptomic studies to infer the relative progression of cells along differentiation trajectories. This approach allows us to study how cells transition from progenitor-like states towards more mature cell types or cell states. Therefore, we performed pseudotime trajectory analysis on ALS brain organoid data to evaluate which embedding best preserves cellular differentiation structure. For each embedding, the data were processed using a standard Seurat workflow. Briefly, the top 3,000 highly variable features were identified, the data were scaled, and dimensionality reduction was performed using PCA. The top 30 principal components were then batch-corrected using Harmony, followed by UMAP visualisation. A nearest-neighbour graph was constructed, and clusters were identified using the FindClusters function at a resolution of 0.4. Cluster re-annotation as performed using canonical cell-type markers from (Szebényi et al. 2021). Pseudotime analysis was performed using the slingshot package. Excitatory neural clusters, identified after re-annotation, were subsetted for trajectory analysis, and intermediate progenitor cells were selected as the starting population. The trajectory analysis results are provided in the main manuscript.

1. <https://www.10xgenomics.com/datasets/3-k-pbm-cs-from-a-healthy-donor-1-standard-1-1-0>
